# Replacing Normalizations with Interval Assumptions Improves the Rigor and Robustness of Differential Expression and Differential Abundance Analyses

**DOI:** 10.1101/2024.10.15.618450

**Authors:** Kyle C. McGovern, Justin D. Silverman

## Abstract

Standard methods for differential expression and differential abundance analysis rely on normalization to address sample-to-sample variation in sequencing depth. However, normalizations imply strict, unrealistic assumptions about the unmeasured scale of biological systems (e.g., microbial load or total cellular transcription). This introduces bias that can lead to false positives and false negatives. To overcome these limitations, we suggest replacing normalizations with interval assumptions. This approach allows researchers to explicitly define plausible lower and upper bounds on the unmeasured biological system’s scale, making these assumptions more realistic, transparent, and flexible than those imposed by traditional normalizations. Compared to recent alternatives like scale models and sensitivity analyses, interval assumptions are easier to use, resulting in potentially reduced false positives and false negatives, and have stronger guarantees of Type-I error control. We make interval assumptions accessible by introducing a modified version of ALDEx2 as a publicly available software package. Through simulations and real data studies, we show these methods can reduce false positives and false negatives compared to normalization-based tools.

## 1 Introduction

Sequence count data (e.g., 16S rRNA-Seq or RNA-Seq data) are ubiquitous in modern biomedical research. These data are often used for Differential Expression or Differential Abundance Analysis (DE/DA). The goal of DE/DA is to determine whether individual microbes (or genes) change in abundance (or expression) between two experimental conditions (e.g., health versus disease). However, limitations of the sequence count measurement process complicate these analyses.

Due to the measurement process, the number of reads sequenced in a sample (i.e., the sequencing depth) is typically independent of the *scale* of the biological system from which that sample was taken (e.g., the total number of microbes in a study participant’s colon) [1, 2, 3]. For this reason, many authors call these data compositional, alluding to the idea that these data only measure the relative amounts of different gene transcripts or microbes within the system, not absolute amounts of transcription or abundance [4]. Yet relative amounts alone are insufficient to perform DE/DA: a microbe could increase in relative abundance but not change, or even decrease, in absolute abundance [5]. To perform DE/DA, we need to know how relative abundances change between conditions and how total abundances (i.e., the system scale) changes between conditions. This latter information can be summarized by a one-dimensional, real-valued quantity *θ*^⊥^ called the *Log-Fold-Change (LFC) in scales* [5, 6, 7]. The core problem is that the observed data provides little to no information about *θ*^⊥^, and therefore modeling assumptions or external measurements of the scale (e.g., flow cytometry) are required.

Until recently, normalizations were the standard approach to addressing the arbitrary scale of sequence count data. Common tools like ALDEx2 [8], DESeq2 [9], and limma [10] use data or parameter normalization to address this scale problem. It is well known that the choice of normalization can dominate study conclusions [4, 5, 11, 12]. We recently showed that normalizations make strong, often implicit, assumptions about scale: they assume that *θ*^⊥^ can be calculated from the observed data without error (Fig. 1A). For example, the common Total Sum Scaling (TSS) normalization corresponds to the unrealistic assumption that there is no change in scale between conditions (*θ*^⊥^ = 0) [7]. Even slight errors in these assumptions can lead to elevated rates of false positives and false negatives in DE/DA. For example, in [6], we showed simulated and real examples where standard normalization-based tools displayed false positive rates above 70% due to such errors. Even more troubling, these error rates typically increase with increasing sample size due to a bizarre statistical phenomenon called *unacknowledged bias*: in failing to acknowledge bias caused by erroneous assumptions about *θ*^⊥^, existing methods become increasingly confident in incorrect results with increasing sample size [5].

**Figure 1:**
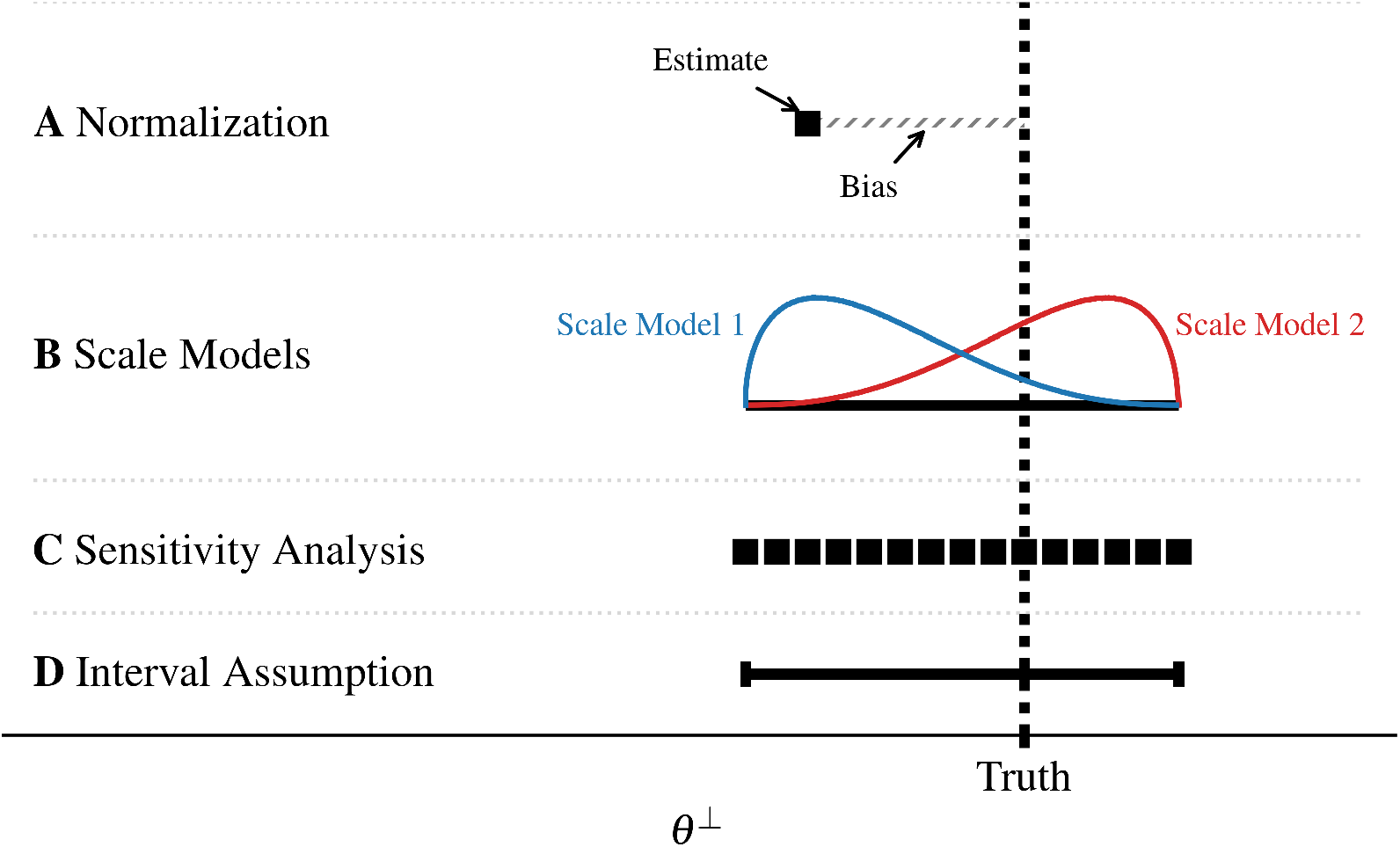
Methods differ in their assumptions about the unmeasured biological system’s scale in DE/DA. A conceptual overview of different approaches in DE/DA for handling the unmeasured LFC in scales: θ^⊥^ (X-axis) is presented visually. The true LFC in scales is marked by a vertical dotted line labeled “Truth”. **A**. Normalization are prevalent; they attempt to estimate the correct θ^⊥^ from the observed data (black point). However, these estimates are inevitably biased (gray hashed lines). **B**. Bayesian scale models define a probability distribution over a range of θ^⊥^ (black line). Of the two example scale models, Scale Model 2 (red) assigns more probability near the truth than Scale Model 1 (blue); thus, an analysis using Scale Model 2 is expected to return fewer false positives. **C**. In a sensitivity analysis, DE/DA is performed repeatedly for distinct values of θ^⊥^ (black points) to identify entities sensitive to the choice of θ^⊥^. **D**. Interval assumptions, introduced here, are defined by lower and upper bounds on θ^⊥^. Robust inference of LFCs and p-values only requires the interval to cover the true θ^⊥^, as depicted.

Recently, we showed that *scale models* could replace normalizations. Scale models are a type of Bayesian prior distribution that represents uncertainty in the unmeasured scale of the system (a distribution *p*(*θ*^⊥^), Fig. 1B) [5, 6]. In a sense, scale models can represent uncertainty in the choice of normalization. Moreover, they can incorporate external scale measurements (e.g., cell concentrations measured by flow cytometry) [6]. By integrating scale models into the popular ALDEx2 software package, we showed that scale model-based methods could dramatically decrease false positive and false negative rates compared to normalization-based methods [6]. Our findings have been replicated on independent studies [13]. However, scale models are not without their limitations. It can be challenging to specify scale models. Even if two researchers agree on a set of biologically plausible values of *θ*^⊥^, they may disagree on which values within that set are more likely. That is, scale models require analysts to choose not only the plausible set but also the distribution over that set (e.g., Gaussian versus uniform) – different distributions can lead to different results.

As an alternative to scale model-based methods, we have also developed sensitivity analyses that take a different approach to addressing the limitations of normalizations [7]. By repeating analyses over different values of *θ*^⊥^, researchers can identify unreliable results that are sensitive to small changes in assumed values of *θ*^⊥^ (Fig. 1C). Sensitivity analyses are simpler than scale models in that they only require researchers to choose a biologically plausible set of *θ*^⊥^. However, sensitivity analyses lack the more familiar statistical constructs of scale models, such as confidence intervals or p-values. While sensitivity analyses can be useful, we expect some researchers will want alternative methods.

This article introduces an alternative to scale model-based and sensitivity analyses for DE/DA. We develop a statistical framework based on interval assumptions of the form 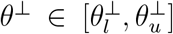 (Fig. 1D). Like scale model-based methods, our interval assumption-based methods provide familiar statistical constructs such as confidence intervals and p-values while maintaining the simplicity of sensitivity analyses. Despite their comparative simplicity, we show that interval assumption-based analyses enjoy many of the same benefits as scale modelbased analyses. Namely, reduced false positive rates compared to equivalent normalization-based analyses and reduced false negative rates when biological prior knowledge or external measurements of scale are incorporated.

### 1.1 A Brief Review of Notation and Concepts from Scale Reliant Inference (SRI)

This article adds to the growing field of *Scale Reliant Inference*(SRI) [5]. In brief, SRI studies the estimation of quantities (e.g., log-fold-changes in DE/DA) that cannot be uniquely identified due to the arbitrary scale of the observed data. A more formal definition of SRI and a comparison to the field of compositional data analysis can be found in Nixon et al. [5]. Consider a study of the gut microbiome where we observed a sequence count dataset *Y* which is a *D ×N* matrix with elements *Y*_*dn*_ representing the number of sequenced reads that map to taxon *d* in sample *n*. Suppose half of these samples were obtained from participants with a disease of interest *x*_*n*_ = 1 and half from healthy controls *x*_*n*_ = 0. The goal of differential abundance analysis is to identify which, if any, of these *D* taxa changed in abundance between the two conditions.

In SRI, we view the observed data *Y* as an imperfect measurement of the underlying biological system *W*. Both *Y* and *W* are *D × N* matrices, but the elements *W*_*dn*_ represent the true, rather than measured, abundance of taxon *d* in the microbial community from which sample *n* was obtained. The system can be uniquely described by its composition *W*^∥^and its scale *W*^⊥^, defined as:

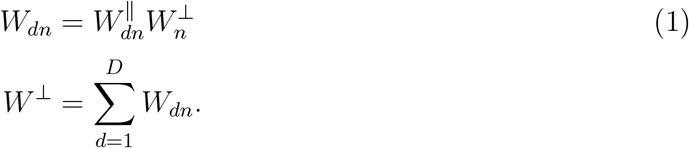

Here *W*^⊥^ is a *D*-length vector with positive elements representing the total amounts (e.g., total microbial load), while *W*^∥^ is a *D×N* matrix of proportional abundances. *Y* is imperfect in at least two ways. First, due to measurement noise, there is some uncertainty in estimating the system composition *W*^∥^ from *Y*. Second, unlike the system composition *W*^∥^, *Y* provides little information about the system scale *W*^⊥^. While the first imperfection–related to the composition–is a standard statistical problem addressed by multiple available methods [8, 14], the second–related to the scale–is less familiar and more troubling.

The lack of information about *W*^⊥^ in the observed data represents a fundamental impediment to DE/DA. DE/DA can be defined as a problem of estimating the Log-Fold-Change (LFC) of each entity *d*:

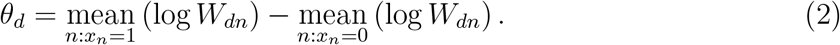

Throughout this work, *log* refers to logarithm in base 2. Using Equations (1) and (2) we can show that DE/DA (i.e., LFC estimation) requires knowledge of the system scale *W*^⊥^:

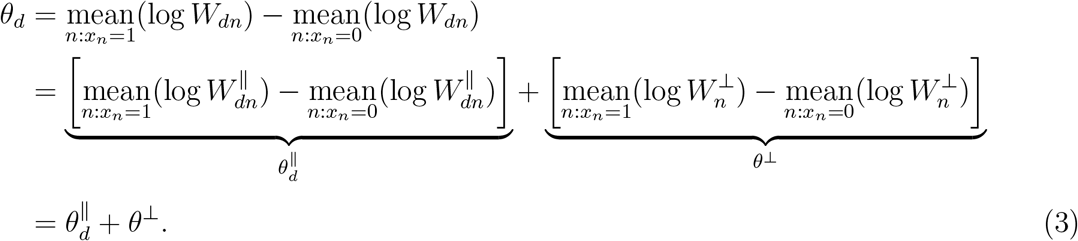

We call 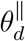 the LFC in composition as it describes how the relative abundance of the *d*-th taxon changes between conditions and *θ*^⊥^ the LFC in scale as it describes how the scale (e.g., total microbial load) changes between conditions. Since sequence count data lack information about *W*^⊥^, these data cannot be used to meaningfully bound *θ*^⊥^ within the range (−∞, +∞). Using Equation (3), it follows that sequence count data alone is insufficient for DE/DA as it cannot bound *θ*_*d*_ ∈ (−∞, +∞). DE/DA cannot be performed using sequence count data alone; it requires external information about the LFC of scales (*θ*^⊥^).

## 2 Results

### 2.1 Interval Assumptions for DE/DA

DE/DA analysis requires external information about the LFC in scales (*θ*^⊥^). This information can be provided by external measurements (e.g., flow cytometry), prior knowledge, or biological plausibility arguments. A flexible and transparent way to state those assumptions or incorporate those measurements is through an interval assumption of the form 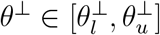.For concreteness, we provide four examples.

**Example 1**. In a microbiome study investigating the effects of antibiotic treatment, researchers may believe that total microbial load is typically lower in patients receiving treatment compared to matched controls. In these cases, an interval of the form 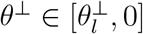 may be appropriate.

**Example 2**. In a microbiome study, researchers might use flow cytometry to measure the total microbial concentration *z*_*n*_ for each sample *n* ∈ *{*1, …, *N}*. They might estimate 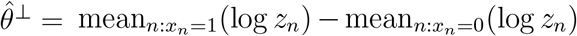 with a standard error *σ*. To account for uncertainty, they might choose an interval 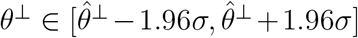 which approximately corresponds to a 95% confidence interval for *θ*^⊥^ based on the central limit theorem.

**Example 3**. In a gene expression study, a researcher who typically uses Centered LogRatio (CLR) normalization for differential expression might use an interval assumption as a more rigorous alternative. The CLR normalization corresponds to the assumption that 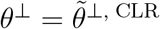 where

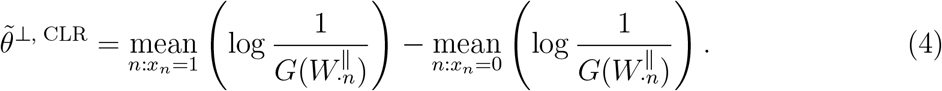

To account for uncertainty in this assumption, this researcher might use an interval assumption of the form 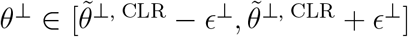 where *ϵ*^⊥^ *>* 0 represents potential error in the estimate.

**Example 4**. In a gene expression study, suppose researchers have a set of known housekeeper genes *H* ⊂ *{*1, …, *D}* that are believed not to change between conditions. In such cases, researchers might use an interval assumption of the form 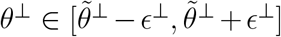 where 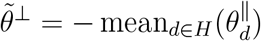 and *ϵ*^⊥^ *>* 0 represents potential error in this assumption.

In the next section we introduce how these interval assumptions share a common form. In later sections, we demonstrate these four types of assumptions though simulation and real data studies.

### 2.2 Interval Assumptions as Generalized Normalizations

All of the interval assumptions in the four examples above share a common form. Many normalization, such as the CLR, can be expressed as functions of the system composition:

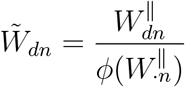

where we have introduced a 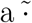 to emphasize 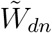 is a quantity derived from normalization. For example, in CLR normalization *ϕ* is the geometric mean function (*G*). With this notation, the interval assumptions discussed previously can be written as

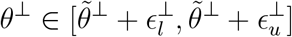

where 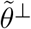 denotes the log-fold-change in scales implied by a particular normalization:

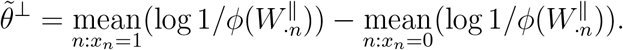

Using this notation, Examples 1 and 2 both illustrate interval assumptions with the trivial normalization: 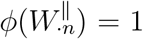,which implies that 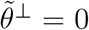.In the first example, 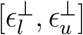 equals 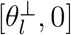.In the second, it equals 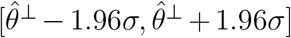.To avoid potential confusion, note 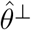 denotes a fixed constant dependent on the flow cytometry data whereas 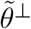 denotes a normalization-based estimate which is a function of the composition *W*^∥^. In the third example, *ϕ* = *G*, and 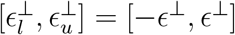.In the fourth example, 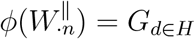 where *G*_*d*∈*H*_is the geometric mean applied only to those genes in the set *H* and 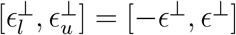.

In summary, each of these four intervals represents potential error 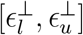 around a normalization-based scale assumption 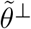,hence we refer to all four as *generalized normalizations*.

### 2.3 From Interval Assumptions to Interval Null Hypotheses

To enable interval assumption-based analyses, we integrate interval assumptions into a hypothesis testing framework. While our approach can be generalized, we assume researchers performing DE/DA are interested in testing the null hypothesis *H*_0_ : *θ*_*d*_ = 0 for each taxon or gene *d*. Using Eq (3), this null hypothesis can be expressed equivalently as 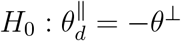.This null hypothesis cannot be tested directly since *θ*^⊥^ is unknown. However, replacing *θ*^⊥^ with an interval assumption of the form 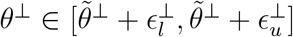.and rearranging the result gives an interval null hypothesis:

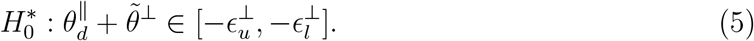

For a given interval assumption, we refer to 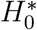 as *the interval null hypothesis*. The superscript ∗ emphasizes that the interval null is not the same as the original null *H*_0_.

We advocate for testing the interval 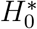 as a proxy for the original null *H*_0_. If a test rejects the interval null hypothesis, it implies we should also reject the original null. This approach is supported by the following key result (proof provided in Supplementary Text 1). If a test *T* (*Y*) controls the false positive rate for the interval null hypothesis, it will also control the false positive rate for the original null hypothesis, provided the interval assumption is correct. Moreover, if the test has non-zero power under the interval alternative 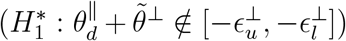, it will also have non-zero power when used as a proxy. In short, so long as the interval assumption is valid, a good test for 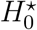 will also be a good test for *H*_0_.

### 2.4 Testing Interval Null Hypotheses and Computing Confidence Intervals

The key to developing tests for these interval null hypotheses is recognizing that the interval null statistic 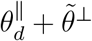 is equivalent to the difference in means of log-transformed, normalized relative abundances:

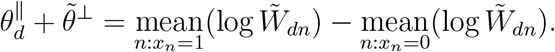

A derivation of this equivalence is provided in Supplementary Text 1. It follows that, when interval assumptions are generalized normalizations, the interval null hypothesis can be tested using standard methods like t-tests or non-parametric analogues. These standard methods can be applied to the log-transformed normalized abundances 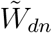,testing whether the difference in means exists within an 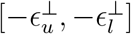.Note that this is a test of a twosided interval null hypothesis, which differs from the more common tests of one-sided nulls or two-sided point nulls.

Readers may be familiar with one-sided tests like the t-test or Wilcoxon test for differencein-means statistics. These tests can be adapted to construct a two-sided interval null test. Let *p*_−_ be the p-value from a one-sided test of 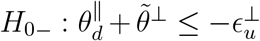,and *p*_+_ the p-value from a test of 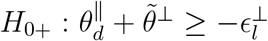. A two-sided interval null test for 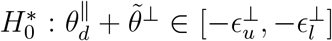 can then be constructed using the composite p-value:

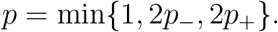

When *p*_+_ and *p*_−_ are calculated using a one-sided t-test, we call this procedure the *Composite T-Test* (CTT).

The CTT is simple, intuitive, and can be extended to other one-sided tests, such as the Wilcoxon test. Additionally, it has connections to sensitivity analysis, as discussed in Supplementary Text 1. However, more powerful tests for interval nulls exist. In *Methods*, we describe the *Generalized T-Test* (GTT), a single, direct test of the interval null that, while less flexible, offers greater power. Furthermore, both the CTT and GTT can be used to construct confidence intervals, as detailed in Supplementary Text 1.

### 2.5 Interval Null Testing in ALDEx2

A challenge in DE/DA is that the composition *W*^∥^ is not directly observed, which prevents calculation of the LFC in the composition 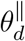 required for testing the interval null hypothesis in Eq (5). Instead, given observed sequence count data *Y*, we have uncertainty in *W*^∥^ that adds to our uncertainty in 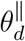.To address this, we take the sampling approach used in ALDEx2 [8] and recently refined when ALDEx2 was updated to incorporate scale models [6]. Given observed sequence count data *Y*, this approach can be summarized in three key steps (detailed in *Methods*). First, we use a Multinomial-Dirichlet bootstrap to account for uncertainty in *W*^∥^ due to small or zero sequence counts [8, 15, 16]. Second, we normalize each bootstrapped sample *W*^∥^ and, for each entity *d*, test the corresponding interval null hypothesis. Finally, for each entity *d*, the resultant p-values are combined into a summary p-value as described in *Methods*.

Coupled with this article, we have released a variant of ALDEx2, which implements the CTT or GTT with any user-specified interval assumption. See *Code Availability* for details.

### 2.6 Interval Assumptions can Dramatically Decrease False Discovery Rates Compared to Normalization

We compared our interval-assumption-based ALDEx2 approach to existing normalization-based methods using the *SparseDOSSA2* simulation framework [17]. SparseDOSSA2 was chosen because its simulation model differs from those of the DE/DA models considered here. SparseDOSSA2 was trained on real metagenomic fecal data to simulate sequence count data matrices *Y* with realistic sparsity, sample-to-sample variance, and microbe-microbe correlation structure. We simulated *D* = 100 microbes with sample sizes in the range *N* ∈ [10, 300] with half the samples treated with a mild selective growth factor and the other half untreated (see *Methods*). Of the 100 microbes, 65 were simulated as increasing in abundance (positive LFC) and 35 unchanging (zero LFC). The distribution of LFCs is shown in Supplementary Figure 1.

We analyzed these data with our variant of ALDEx2 modified to incorporate interval assumptions and the CTT and GTT tests. We refer to these as ALDEx2-CTT and ALDEx2-GTT, respectively. Given that the true simulated LFC in scales was *θ*^⊥^ = 1.73, we tested one correct interval assumption: *θ*^⊥^ ∈ [0, 1.75] and one misspecified interval assumption: *θ*^⊥^ ∈ [0, 0.5]. The correct interval assesses the models’ ability to control false positives, while the misspecified interval assesses the models’ performance under model misspecification. In later sections, we will demonstrate how to define interval assumptions in practice from prior biological knowledge when the true *θ*^⊥^ is unknown.

In Fig. 2, we compared ALDEx2-CTT and ALDEx2-GTT (with both interval assumptions) against four normalization-based methods: ALDEx2 (with CLR normalization) [8], DESeq2 [9], limma [10], and edgeR [18], using False Discovery Rate (FDR) and power across different sample sizes. As expected, given Theorem 1 in Supplementary Text 1, ALDEx2-CTT and ALDEx2-GTT maintained FDR control at the target level of 0.05 when using the correct interval assumption *θ*^⊥^ ∈ [0, 1.75] (Fig. 2A). In contrast, all normalization-based methods failed to control the FDR at almost all sample sizes. Even with the misspecified interval assumption *θ*^⊥^ ∈ [0, 0.5], our methods outperformed all normalization-based approaches in FDR control. While FDR increases with sample size under misspecification, it does so much slower than with the four normalization-based methods. Critically, since our method is a variation of ALDEx2, our superior FDR control compared to ALDEx2 is solely due to replacing CLR normalization with an interval assumption. However, our method’s superior FDR control came with a moderate reduction in power compared to most normalization-based methods. At a sample size of 300, ALDEx2-GTT with the correctly specified interval had a power of 0.47, compared to 0.68 for the most powerful normalization-based method, limma (Fig. 2B). While limma was more powerful, that power came at a steep cost to the trustworthiness of its results: at a sample size of 300, one in two microbes identified as differentially abundant by limma were false positives. In contrast, at the same sample size, only one in 3000 microbes identified as differentially abundant by ALDEx2-GTT with the correct interval assumption were false positives.

**Figure 2:**
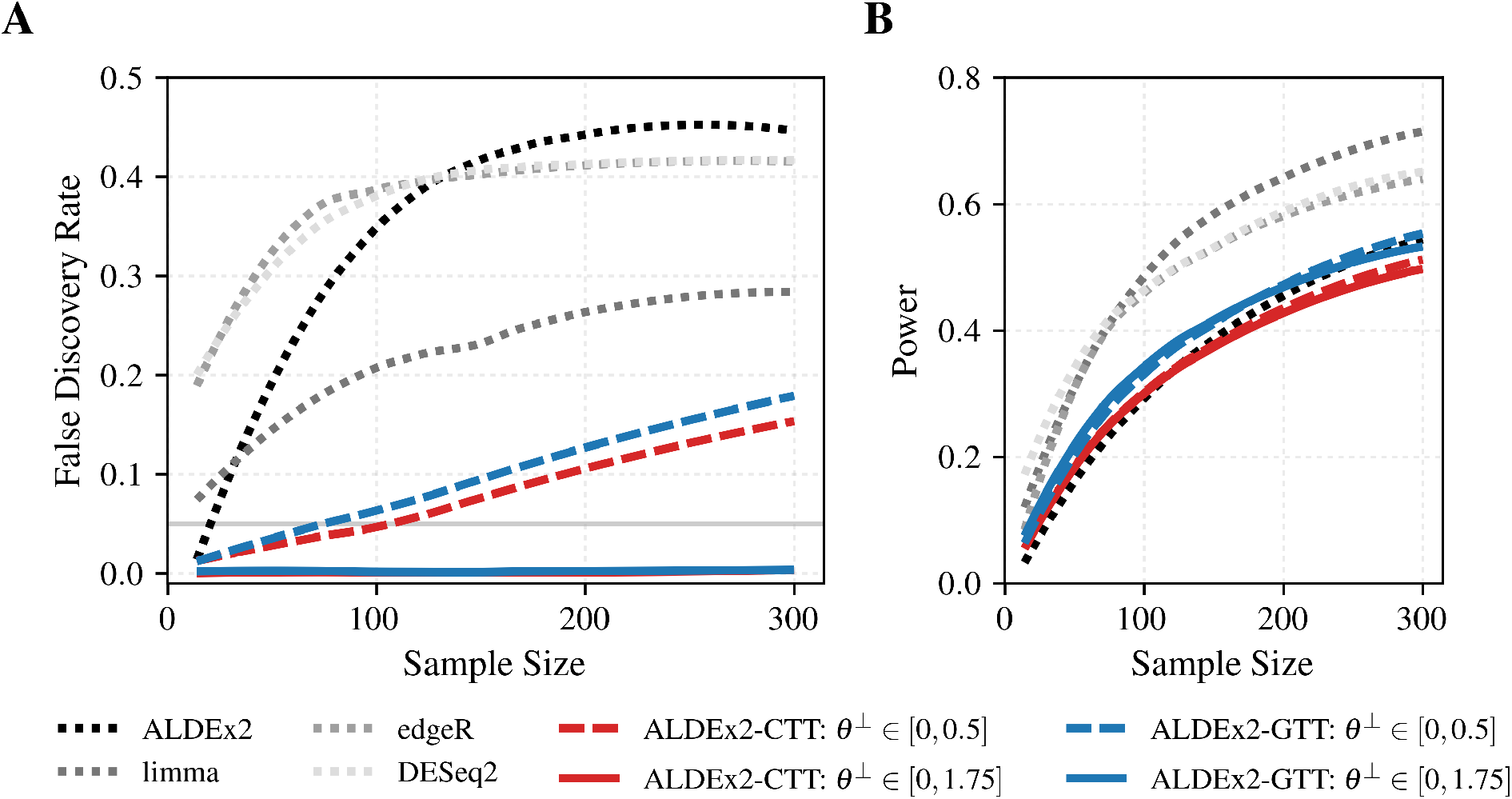
Interval assumptions enable control of the false discovery rate, unlike normalization. Four normalization-based methods, DESeq2, limma, edgeR, and ALDEx2, were benchmarked against the interval-assumption-based methods ALDEx2-CTT and ALDEx2-GTT using simulations of microbiomes in a control vs. mild selective growth factor condition. The true simulated LFC in scales was θ^⊥^ = 1.72, thus the interval assumption θ^⊥^ ∈ [0, 1.75] is correct while the interval assumption θ^⊥^ ∈ [0, 0.5] is incorrect. **A**. The average false discovery rate was computed over 100 independent simulations at sample sizes in the range [10, 300] for each DE/DA method. Only ALDEx2-CTT and ALDEx2-GTT with the correct interval assumption (θ^⊥^ ∈ [0, 1.75]) controlled the false discovery rate at or below the desired level 0.05 (gray solid line). **B**. The average power was computed for each simulation as well.

Finally, we performed an additional, simple simulation study (with the LFC in Supplementary Figure 2A) to illustrate how interval assumptions are simpler than scale models. We created three scale models: a uniformly distributed, a left-skewed, and a right-skewed scale model (Supplementary Figure 2B; see *Methods*). Each of the three scale models was defined over the same support: *p*(*θ*^⊥^) *>* 0 if and only if *θ*^⊥^ ∈ [−1.25, 1.25]. We compared ALDEx2 with these three scale models to ALDEx2-CTT and ALDEx2-GTT using the interval assumption *θ*^⊥^ ∈ [−1.25, 1.25], which matches the support of the scale models. All methods, therefore, covered the true simulated *θ*^⊥^ = 1.22. As expected, both ALDEx2-CTT and ALDEx2-GTT controlled the FDR at all sample sizes tested (i.e., *N* ∈ [0, 200]). In comparison, only ALDEx2 with the right-skewed scale model controlled FDR. In contrast, ALDEx2 with the left-skewed scale model had an even higher FDR than the original ALDEx2 (Supplementary Fig 2C). These results highlight that scale models require not only an analyst to choose the support of the scale model (analogous to what is required to use interval assumptions) but also to specify the probability of different values within that support. Still, our results should not be taken as an admonishment of scale models. The right-skewed scale model is more powerful than our interval-based methods, which are more conservative (Supplementary Figure 2D).

### 2.7 Interval Assumptions Can Decrease False Positives and False Negatives in Real Data

We reanalyzed an oral microbiome study of tooth-brushing, which collected an equal number of unstimulated saliva samples before and after brushing (*N* = 32) [19]. Both 16S rRNA-seq and flow cytometry were performed to quantify the relative abundances of the genera (i.e., the composition) and total microbial load (i.e., the scale), respectively.

The flow cytometry data provided a gold standard scale measurement against which DE/DA tools could be compared. Using a mixed effects model (see *Methods*), we calculated the 95% Confidence Interval (CI) as *θ*^⊥^ ∈ [−1.89, −0.43], reflecting a −73% to −26% change in microbial load after brushing. We applied ALDEx2-GTT to the sequence count data using this CI as an interval assumption. We take the results of ALDEx2-GTT to be a gold standard for identifying true/false positives and negatives. Our gold-standard analysis identified the genera *Streptococcus* and *Haemophilus* as decreased in abundance after brushing (Fig. 3A). These results are consistent with our knowledge of the biogeography of plaque biofilm: *Haemophilus* and *Streptococcus* are frequently found in close proximity in the periphery of plaque biofilms, where they can be removed easily by brushing [20].

**Figure 3:**
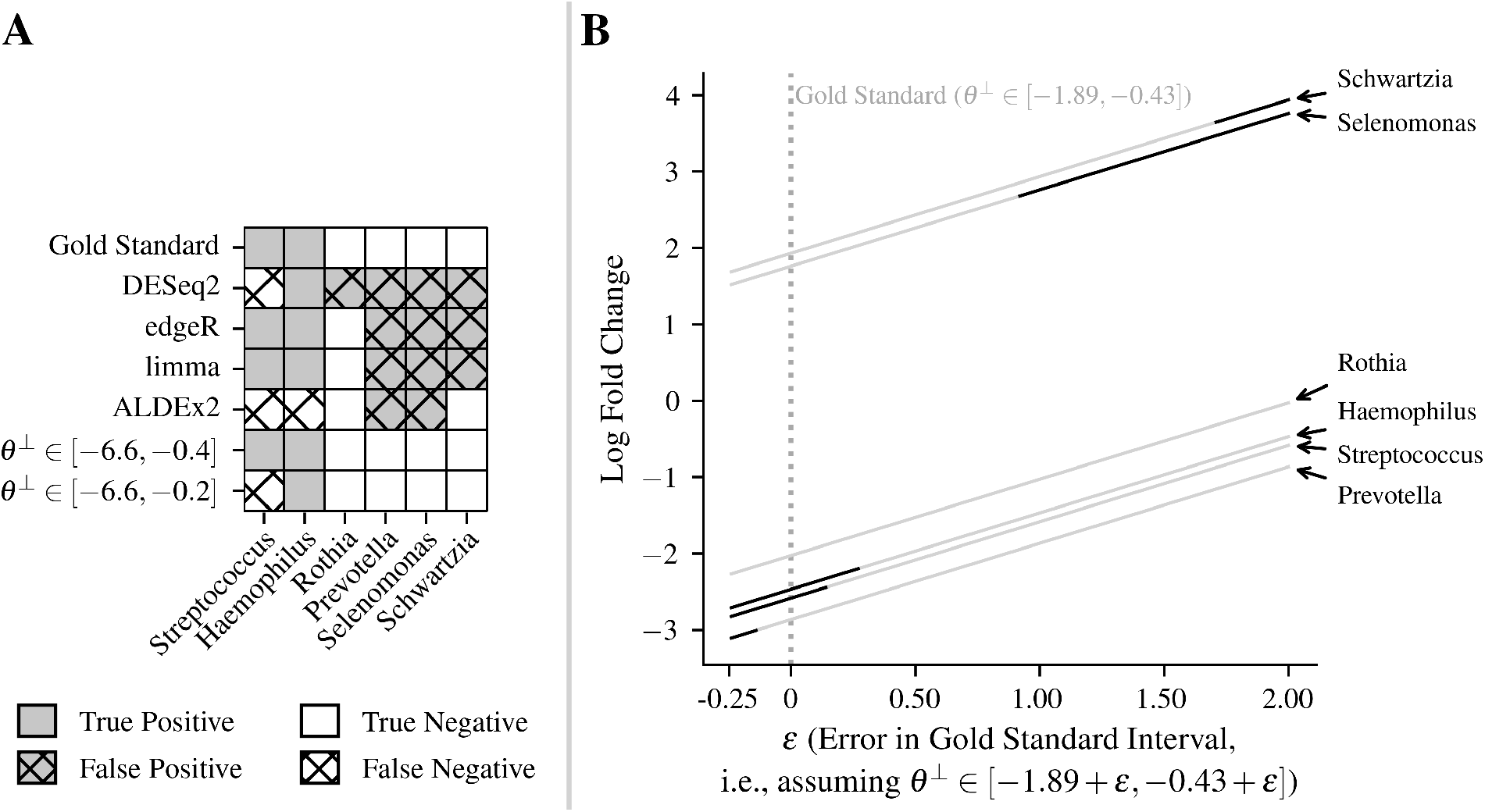
Interval assumptions based on weak prior biological knowledge can reduce false positives and false negatives. **A**. Comparison of true/false positives and negatives across different DE/DA tools applied to the Morton et al. tooth-brushing dataset. Only genera identified as differentially abundant by at least one tool are shown. The gold standard was calculated by ALDEx2-GTT and used the interval assumption θ^⊥^ ∈ [−1.89, −0.43] based on a 95% confidence interval from flow cytometry measurements of oral microbial load. Results from ALDEx2-GTT using the interval assumptions θ^⊥^ ∈ [−6.6, −0.4] and θ^⊥^ ∈ [−6.6, −0.4] are also shown, derived from a literature review of tooth-brushing efficacy. **B**. Sensitivity analysis of ALDEx2-GTT with varying interval assumptions θ^⊥^ ∈ [−1.89 + ϵ, −0.43 + ϵ], where ϵ represents deviation (i.e., error) in the gold standard. When ϵ = 0, this corresponds to the gold standard, as denoted by the gray dotted line. Each solid line represents a genus and is light gray if the Benjamini-Hochberg (BH) adjusted p-value > 0.05 and black if ≤ 0.05.

We compared the gold standard to the same four common normalization-based methods as before (Fig. 3A). All four methods identified *Prevotella* as decreasing and *Selenomonas* as increasing in abundance after brushing, in disagreement with the gold standard. All methods except ALDEx2 identified *Schwartzia* as increasing in abundance after brushing, also disagreeing with the gold standard. *Prevotella* is abundant in plaque biofilms, although it is more common in subgingival plaque where it is difficult to remove by brushing [20]. The increase in *Selenomonas* and *Schwartzia* is unexplained: these genera are not even common contaminants of toothbrushes [21]. To assess the support of these results by the observed data, we performed a sensitivity analysis by recalculating the gold standard with interval assumptions *θ*^⊥^ ∈ [−1.89 + *ϵ*, −0.43 + *ϵ*] where *ϵ* ranged from −0.25 to 2 (Fig. 3B). *Prevotella* was differentially abundant only when *ϵ* ≤ −0.3; with *ϵ* = −0.3 corresponding to the interval assumption *θ*^⊥^ ∈ [−2.19, −0.73]. We used the sampling distribution of *θ*^⊥^ from the mixed-effects model to quantify the degree to which the flow cytometry data support this interval (see *Methods*). This interval covers 87% of the sampling distribution, indicating that the conclusion that *Prevotella* is differentially abundant could be consistent with the flow cytometry data. In contrast, *Selenomonas* and *Schwartzia* were significant only with *ϵ* ≥ 1.3 and *ϵ* ≥ 2.0, respectively. Their corresponding intervals (*θ*^⊥^ ∈ [−0.59, 0.87] and *θ*^⊥^ ∈ [0.11, 1.57]) covered only 6.2% and 0.02% of the sampling distribution, indicating inconsistency between their results and the flow cytometry data. Overall, the four normalization-based methods (ALDEx2, edgeR, limma, and DESeq2) exhibited multiple false positives and some false negatives in this dataset.

Finally, we explored interval assumptions without flow cytometry data, relying solely on a literature review. We found a systematic review that estimated a 99% reduction in plaque index scores from *in vitro* tooth-brushing studies and a 25% reduction from *in vivo* studies [22]. To be conservative, we considered all possible reductions in microbial load between these two values, corresponding to the interval assumption *θ*^⊥^ ∈ [−6.6, −0.4]. Remarkably, using an interval informed only by literature review resulted in perfect agreement with the gold standard (Fig. 3A). Further literature review identified an *in vitro* study that reported a smaller 15% reduction in plaque biofilm as measured by Colony Forming Units (CFUs) [23]. Expanding our interval to *θ*^⊥^ ∈ [−6.6, −0.2] to incorporate this more conservative estimate yielded only a single false negative (Streptococcus; Fig. 3A). Even with this expanded interval, our methods were superior to normalization-based methods in terms of the overall number of inferential errors (false positives plus false negatives). Overall, these results highlight that weak prior knowledge about scale can be sufficient to define interval assumptions and ensure robust DE/DA inference.

### 2.8 Interval Assumptions Can Address Error When Normalizing to Housekeeping Genes

Researchers often normalize gene expression data using housekeeping genes. This approach assumes that the putative housekeeper genes’ expression is truly constant [24]. This approach can introduce bias if the gene expression of those genes is not constant. Previously [13] showed that scale models could be used to account for errors in this constant expression assumption, providing more robust inference. Here we demonstrate interval assumptions for this task.

We demonstrate this with an RNA-seq dataset from The Cancer Genome Atlas (TCGA), comparing the expression of *D* = 18992 genes in normal renal tissue (*n* = 40) versus Clear Cell Renal Cell Carcinoma (CCRCC) tissue (*n* = 40) [25]. We first used ALDEx2 for analysis, normalizing with *GAPDH* as a housekeeping gene (see *Methods*). This approach assumes *θ*_GAPDH_ = 0, making it only reliable if *GAPDH* does not change between experimental conditions. However, we intentionally chose *GAPDH* as a housekeeping gene because it is typically over-expressed in carcinomas and linked to reduced patient survival [26, 27]. As expected, normalization with *GAPDH* leads to likely false positives. For instance, *PPIA* and *TBP* are among the 15877 genes called differentially expressed (see Supplementary Table 1), even though these genes have been confirmed by rt-qPCR to have stable expression between healthy and CCRCC tissues [28].

To address potential error in the assumption *θ*_GAPDH_ = 0, we used ALDEx2-GTT with an interval assumption. Using the fact Eq. (3) implies *θ*_*GAPDH*_ = 0 is equivalent to 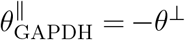, we defined an interval assumption as:

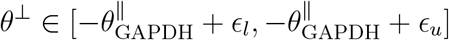

with *ϵ*_*l*_ = 0 and *ϵ*_*u*_= 2.5, based on prior rt-qPCR and micro-array analyses showing the LFC of *GAPDH* to be as high as 2.4 in some tumors [29]. Following Eq. (5), this led to the interval null hypothesis:

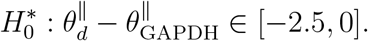

ALDEx2-GTT testing of this hypothesis identified 1500 differentially expressed genes (see Supplementary Table 2), including likely true positives *NNMT* and *VEGFA*, which are known to be over-expressed in CCRCC and linked to reduced patient survival [26, 27]. Critically, ALDEx2-GTT did not label the likely false positives *PPIA* and *TBP* from the previous analysis as significant. However, this approach may require defining a broad interval, which can reduce statistical power. For instance, the genes *GDF15* and *PPARA*, (known to be down-regulated in CCRCC) were identified as differentially expressed by ALDEx2 but not ALDEx2-GTT [30, 31].

Overall, these results align with the previous simulation analyses. Our approach may miss true positives, such as the *GDF15* and *PPARA* genes. Yet those genes that are identified as differentially expressed by ALDEx2-GTT, like *NNMT* and *VEGFA*, are well-supported by the literature. In contrast, while ALDEx2 identified all these genes, it critically compromised the reliability of its results. A researcher using ALDEx2 with *GAPDH* as a housekeeping genes might erroneously conclude the genes *PPIA* and *TBP* are differentially expressed, despite the available literature suggesting otherwise.

## 3 Discussion

Common DE/DA tools typically rely on normalization to overcome the arbitrary scale of sequence count data. However, normalization methods imply strict model assumptions, which can introduce bias and confound scientific conclusions. We have previously proposed scale models and sensitivity analyses to address these issues. While useful, scale models can be challenging to specify, and sensitivity analyses lack familiar statistical constructs like p-values and confidence intervals. In this work, we proposed interval assumptions and developed a framework for testing interval null hypotheses based on these assumptions. We have made these methods publicly available as a software package. Simulation and real data studies showed that our approach can drastically reduce false positive rates compared to normalization-based methods. Additionally, our method maintains lower false positive rates even with moderate misspecification of interval assumptions. Moreover, we showed that interval assumptions informed by prior knowledge or external scale measurements can even reduce false negative rates compared to existing methods. Overall, this work introduces a novel and flexible approach to DE/DA analysis that improves robustness by addressing potential errors in modeling assumptions about scale.

We expect potential confusion regarding our claim of Type-I error control (in Theorem 1 in Supplementary Text 1) compared to the results of Nixon et al. [5]. In that work, we proved that Type-I error control with non-trivial power is impossible when performing DE/DA from data with arbitrary scale. Yet, this article has developed a framework from DE/DA that can control Type-I error with nontrivial power. The distinction is that the Type-I error control we discuss here is weaker than that discussed in the prior article. There we discussed classical Frequentist Type-I error control defined as a worst-case guarantee over all possible models (e.g., all possible scales *θ*^⊥^ ∈ (−∞, ∞)). This article discusses a weaker form of Type-I error control that is only guaranteed conditioned on an interval assumption 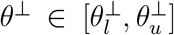 being true. Still, our methods are useful. Consider that assumptions are inevitable. Normalizations-based methods also make assumptions yet those assumptions are often much stronger (e.g., *θ*^⊥^ = 0). In contrast, our interval assumptions are weaker (e.g., *θ*^⊥^ ∈ (−*ϵ, ϵ*)) and are therefore more likely to be satisfied in practice.

Our approach is related to, yet distinct from, shape constraints in the econometrics literature. Scale Reliant Inference (SRI) is formulated as a problem of estimation in partially identified models, where parameters are not uniquely identified due to the lack of scale information in the observed data [5]. In the econometrics literature, partially identified models arise due to *shape constraints*: geometric restrictions such as monotonicity between treatment and effect that ensure more plausible models [32]. Interval assumptions are shape restrictions on *θ*^⊥^. However, to our knowledge, we are the first, even within the econometrics literature, to convert shape constraints into interval null hypotheses. A key advantage of our approach is that we can use off-the-shelf tools to test those hypotheses. Overall, we expect future works adapting ideas from the econometrics literature on shape restrictions to the study of sequence count data to be impactful.

Our approach also relates to commonly used LFC thresholds in DE/DA analyses [12, 33, 34]. In brief, these analyses typically put thresholds on both p-values (e.g., *p <* 0.05) and the estimated LFC (e.g., |*θ*_*d*_| ≥ 1) as requirements for a gene or microbe to be called differentially expressed or abundant. Our method is similar: if we assume that *θ*^⊥^ ∈ (−*ϵ, ϵ*) we are requiring that 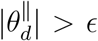 in order to reject the null hypothesis. Still, our approach is distinct for at least two reasons. First, our approach accounts for estimation uncertainty in 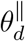,whereas LFC thresholds do not. Second, thresholds on LFC (*θ*_*d*_) depend on often biased modeling assumptions about *θ*^⊥^ (e.g., from normalization), which can make the thresholds themselves biased. In contrast, our approach can place cutoffs directly on 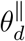.which can be learned from the data, thus avoiding bias.

This article has developed a method for DE/DA that addresses two imperfections of sequence count data: (1) limited sequencing depth which results in uncertainty in the LFC in composition 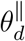,and (2) the lack of scale information in the observed data which leads to substantial uncertainty in the LFC in scales *θ*^⊥^. However, this work does not address other possible imperfections (e.g., PCR bias or batch effects) that may bias our estimates of 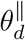.Such bias could be addressed by interval assumptions, similar to how scale was addressed here. For concreteness, suppose compositional estimates were biased such that the true value 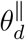 was related to our estimate 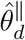 by some systematic error 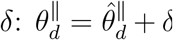.Using prior literature (e.g., [35, 36, 37]), we could specify an interval assumption of the form δ ∈ [δ_*l*_, δ_*u*_]. Functionally, this could be incorporated into our framework alongside scale assumptions of the form 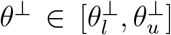 via a combined interval 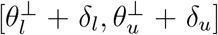,which could be turned into an interval null hypothesis in the same way as we have discussed. Still, this may not be necessary for some sources of bias. For example, we and others have provided theory and experimental results that suggest that DE/DA analyses may be unaffected by PCR bias [35, 36].

Overall, our interval assumption approach complements the scale model approach and sensitivity analyses to create a suite of tools for scale-reliant inference from sequence count data. We guide researchers in navigating these distinct approaches. Researchers familiar with Bayesian modeling might prefer scale models (e.g., [5, 6]) for the superior flexibility and power that comes from the ability to specify different probabilities to different *θ*^⊥^. Still, this flexibility has potential pitfalls: poorly specified scale models that do not accurately reflect prior beliefs can lead to inferential errors. Other researchers may prefer interval assumptions because they are more familiar with Frequentist tools (e.g., calibrated p-values from null hypothesis tests), are simpler to specify, and often have better control of the false positive rate. Finally, sensitivity analyses (e.g., [7]) are a distinct tool. Sensitivity analyses are not part of a formal inferential framework (e.g., they do not test hypotheses or provide distinct estimates of a parameter of interest). Instead, sensitivity analyses should be considered a type of data visualization: one can use sensitivity analyses to visualize which study conclusions are sensitive and which are insensitive to errors in modeling assumptions [7].

## 4 Methods

### 4.1 The Generalized T-Test

We summarize the generalized t-test developed by Mehring [38]. Further details, derivations, and proofs related to Type-I error and power can be found in that article. For two sets of observed data, *x*_1_, …, *x*_*N*_and *y*_1_, …, *y*_*M*_ drawn from two populations where *x*_*n*_ ∼ 𝒩 (*µ*_1_, *σ*^2^) and *y*_*m*_ ∼ 𝒩 (*µ*_2_, *σ*^2^), the GTT tests the interval null hypothesis *H*_0_ : *µ*_Δ_ ∈ [*ψ*_1_, *ψ*_2_] where *µ*_Δ_ = *µ*_2_ − *µ*_1_ and *ψ*_1_ *< ψ*_2_.

Without loss of generality, by using the interval midpoint: 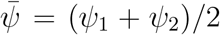,this null hypothesis can be reformulated as 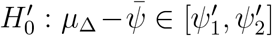 where 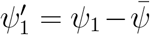 and 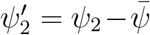.

This redefinition ensures that 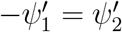 and 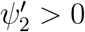.

The GTT p-value for 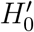 is defined as

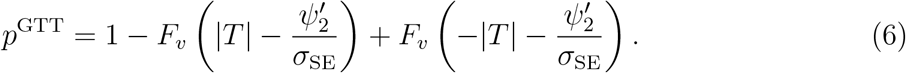

where *T* is the Student’s t-statistic, *σ*_SE_ is the standard error for the difference in means, and *F*_*v*_ is the cumulative density function of the t-distribution with *v* degrees of freedom. *T, σ*_SE_, and *v* are calculated using standard formulas for the two-sample Student’s t-test with equal variances.

### 4.2 Interval Null Hypothesis Testing in DE/DA with a Modified ALDEx2 Model

We developed a method and software tool for interval null hypothesis testing in DE/DA based on a modification to the ALDEx2 framework proposed by Fernandes et al. and recently updated by Nixon et al. [6, 8]. Our method involves three key steps.

First, *S* Monte Carlo samples are drawn from the posterior of *N* independent multinomial-Dirichlet distributions to estimate the composition. For Monte Carlo sample *s* and sample *n*, the posterior sample is:

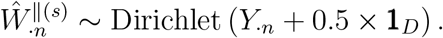

where 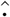 denotes an estimate, *Y*_·*n*_ is the *D*-length vector of observed sequence counts for sample *n*, and **1**_*D*_ denotes a *D*-length column-vector of ones.

Second, we calculate normalized estimates 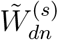 using:

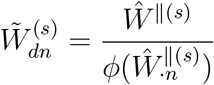

where *ϕ* is the generalized normalization function. For CLR normalization, *ϕ* is the geometric mean (*G*). Our software uses 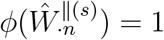 as the default, leading to 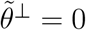 and an interval assumption of the form 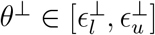.For each gene or taxon *d* we then compute:

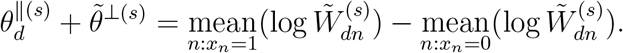

Then we apply the CTT or GTT test of the interval null hypothesis in Eq. (5) using the user-defined bounds 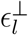 and 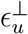 to calculate a 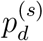 for each Monte Carlo sample *s*.

Standard errors for these tests are computed from the calculated 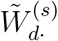.

Finally, the p-values 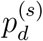 for *s* = 1, …, *S* are summarized into a single p-value *p*_*d*_. Following Nixon et al., summarization was defined to account for changes in the sign of the LFC [6]. 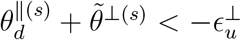,the LFC of *d* must be negative, while if 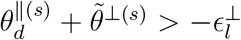 the LFC of *d* must be positive. If 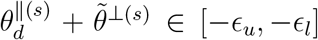,the LFC of *d* may be zero, positive, or negative. The simple averaging procedure 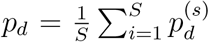 fails to account for changes in the sign of the LFC between Monte Carlo samples. Instead, we first calculate lower and upper p-values: 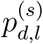 and 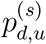, as

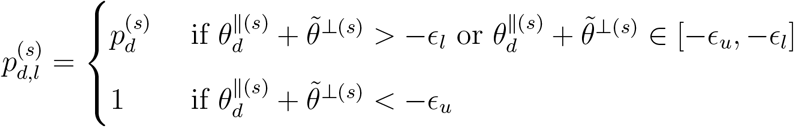

and

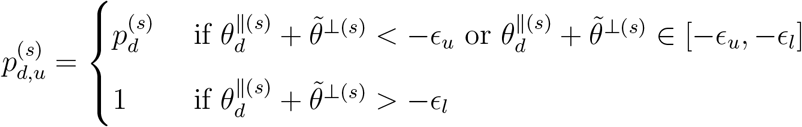

The summary p-value for each entity *d* is then calculated as

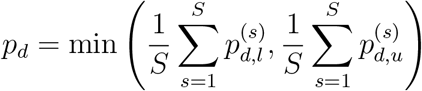

To correct for multiple hypothesis testing, Benjamini-Hochberg adjustment is applied 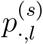and 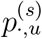 before computing *p*.

### 4.3 Simulation Studies

For both simulation studies and all DE/DA methods, a microbe *d* was deemed significant if its Benjamini-Hochberg (BH) adjusted p-value was below 0.05. ALDEx2 (version 1.35.0 with scale model functionality), ALDEx2-CTT, and ALDEx2-GTT used 300 Monte Carlo samples. For ALDEx2-GTT and ALDEx2-CTT, 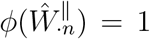 was used 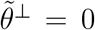.

DESeq2 (version 1.42.1) was applied using the “poscounts” method to estimate size factors, accommodating microbes with many zeros. For limma (version 3.58.1) and edgeR (version 4.0.16), default parameters were applied. Reported power and false discovery rates for all simulations were averages over 100 independent runs.

#### 4.3.1 Mild Growth Factor Simulation

Sequence count data for each sample size *N* between 10 and 300 were simulated using SparseDOSSA2 (version 0.99.2) with the default “stool” model, pre-trained on HMP1-II healthy human stool metagenomic samples [17]. For half the samples under the mild growth factor condition, LFCs were simulated using the abundance spike-in feature. To account for an expected decrease in sparsity with higher abundance, the prevalence spike-in feature was set to half the microbe’s LFC, reducing the number of absolute zeros for microbes with positive LFCs. To further reduce sparsity in the analysis and remove low-signal microbes, microbes with non-zero counts in fewer than 25% of samples were combined into a microbe category labeled “other”. This resulted in *D* = 100 microbes analyzed, including “other”. Sequencing depth per sample was simulated uniformly between 5, 000 and 100, 000.

#### 4.3.2 Simulation Benchmarking of Scale Models

To benchmark ALDEx2-CTT and ALDEx2-GTT against ALDEx2 using scale models, absolute abundances for *D* = 100 microbes were simulated from multivariate normal distributions as

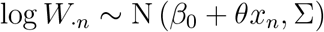

where *x*_*n*_ is a binary covariate indicating case (*x*_*n*_ = 1) or control (*x*_*n*_ = 0), Σ is a covariance matrix simulated from an inverse-Wishart distribution with *D* + 3 = 103 degrees of freedom, and the intercept *β*_0_ is an *D*-length column vector with elements drawn from a standard normal distribution. The absolute abundances *W*_·*n*_ were converted into 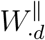 using Eq. (1) and then used to simulate sequence counts *Y*_·*n*_ from a multinomial distribution with a sequencing depth of one million reads.

#### 4.3.3 Scale Models in ALDEx2

ALDEx2 scale models were implemented using the ALDEx2 framework described by Nixon et al. [6] using the *γ* parameter. The parameter *γ* is an *N × S* matrix where *γ*_*n*s_ represents the scale 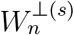 for observed sample *n* and Monte Carlo sample *s*. The scale model was defined as

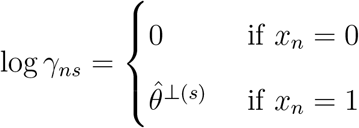

where *x*_*n*_is a binary covariate indicating case (x_*n*_ = 1) or control (x_*n*_ = 0). For each Monte Carlo sample s,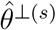 is constant across all samples *n*. In the uniform scale 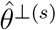 was sampled uniformly from [−1.25, 1.25]. In the skewed scale models, 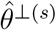 was drawn from four-parameter Beta distributions: Beta(1.25, 8, −1.25, 1.25) for the left-skewed model and Beta(8, 1.25, −1.25, 1.25) for the right-skewed model.

### 4.4 Reanalysis of Teeth-Brushing Oral Microbiome Study

Data analysis using ALDEx2 and ALDEx2-GTT was conducted with 1000 Monte Carlo samples. For DESeq2, limma, and edgeR, default parameters were applied. For ALDEx2-GTT, 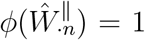 was used implying 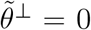.Microbes were considered significant if their Benjamini-Hochberg adjusted p-values were below 0.05.

#### 4.4.1 Preprocessing of Sequence Count Data

A 16S rRNA-seq count matrix (*N* = 32) from 10 study participants, pre-processed according to Morton et al. [19], was obtained from the European Bioinformatics Institute (EBI) database under accession number ERP111447. Sequence counts were aggregated at the genus level for analyses. Genera with non-zero counts in fewer than 12 samples were amalgamated into a single genus labeled “other” to retain the total sequencing depth, resulting in *D* = 38 total genera for subsequent analyses.

#### 4.4.2 Mixed Effects Modeling of Flow Cytometry Data

Flow cytometry cell counts, preprocessed as described by Morton et al., were obtained from the Qiita database (study number 11896) and treated as scale measurements [19]. The dataset included 64 measurements, with two technical replicates per each of the 32 biological samples. These measurements were analyzed on the log scale using a mixed effects model with treatment (before vs. after brushing) as a fixed effect, and random intercepts for study participant and time of day (morning vs. evening). The model, fitted using the R package lme4 (version 1.1-35.5), was specified as

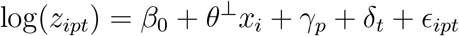

where *z*_ipt_ represents the *i*-th flow cytometry measurement from the *p*-th study participant at time of day *t*. The random effects are 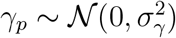 and 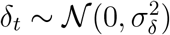 for study participant *p* and time of day *t*, respectively; and *ϵ*_ipt_ ∼ 𝒩 (0, *σ*^2^) is residual error. Here, *β*_0_ is a fixed intercept term and *x*_i_ is a binary covariate indicating if measurement *i* was from before (*x*_i_ = 0) or after (*x*_i_ = 1) brushing.

From this mixed effects model, the estimated scale was 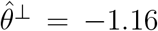 with a standard error 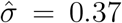.The 95% confidence interval estimated using the normal approximation 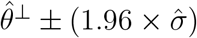 was estimated as [−1.89, −0.43].

#### 4.4.3 Gold Standard Analysis

ALDEx2-GTT computed the gold standard results using the 95% confidence interval from the mixed effects model analysis (*θ*^⊥^ ∈ [−1.89, −0.43]) as the interval assumption. This interval assumption corresponded to testing the interval null hypothesis 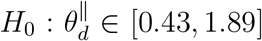 using Eq. (5).

#### 4.4.4 Sensitivity Analysis

For the sensitivity analysis with ALDEx2-GTT, we used the interval assumption *θ*^⊥^ ∈ [−1.89 + *ϵ*, −0.43 + *ϵ*], where *ϵ* ∈ [−0.25, 2] represented error in the gold standard. For each *ϵ*, this corresponded to the interval null hypothesis 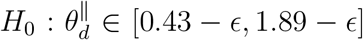.Coverage of the sampling distribution, derived from the mixed effects model as *θ*^⊥^ ∼ 𝒩 (−1.16, 0.37), was calculated for each *ϵ* as

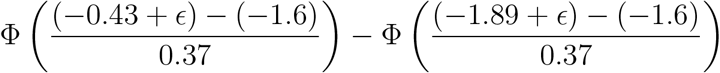

where Φ is the cumulative density function of the normal distribution.

### 4.5 Differential Expression Analyses of Clear Cell Renal Cell Carcinoma Tissues

#### 4.5.1 Preprocessing of Sequence Count Data

Metadata and sequence count data, pre-processed as described by Rahman et al. [25], were obtained from NCBI’s Gene Expression Omnibus (GEO) database (accession numbers GSE62944, GSM1536837, and GSM1697009). We randomly selected 40 samples from both normal and CCRCC tissues, totaling *N* = 80 samples. Genes with non-zero counts in at least 25% of samples were retained, resulting in *D* = 18, 992 of the 23368 genes used for analysis. The remaining 4, 376 genes were combined into an “other” category.

#### 4.5.2 Differential Expression Analyses

For both the ALDEx2 and ALDEx2-GTT analyses, genes were considered differentially expressed if their Benjamini-Hochberg adjusted p-values were below 0.05. Each analysis utilized 250 Monte Carlo samples. ALDEx2 normalization was performed using *GAPDH* as a reference gene by setting the “denom” parameter to *GAPDH* ‘s gene index. Equivalently, for ALDEx2-GTT, the generalized normalization 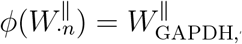 was used, which implied

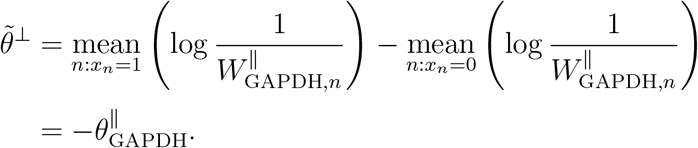

This relationship was then used to derive the interval assumption and interval null hypothesis presented in *Results*.

### 4.6 Data Availability

All data used in this manuscript have been previously published and are publicly available. The data for the oral microbiome analysis is available from the European Bioinformatics Institute (EBI) database under accession number ERP111447 and the Qiita database under study number 11896. The metadata and sequence count data for the Clear Cell Renal Cell Carcinoma analysis are available from NCBI’s Gene Expression Omnibus (GEO) under accession numbers GSE62944, GSM1536837, and GSM1697009.

### 4.7 Code Availability

All code needed to generate the figures, supplementary figures, and supplementary tables are provided at: https://github.com/Silverman-Lab/Interval-Null-DE-DA. The modified ALDEx2 model for testing interval null hypotheses is implemented as an R software package called INDExA and is available at: https://github.com/Silverman-Lab/INDExA.

