## Supplementary Text and Figures for "Replacing Normalizations with Interval Assumptions Improves the Rigor and Robustness of Differential Expression and Differential Abundance Analyses"

### Supplementary Text 1

#### Statistical theory for interval null hypothesis testing in DE/DA

##### 0.1 Notation and Definitions

Here we introduce notation and definitions necessary for this section, following the Scale Reliant Inference (SRI) framework of Nixon et al. [1].

Let $P_{a,b}$ denote a probability model which is a function mapping a parameter space $\mathcal{A} \times \mathcal{B}$ to a set of distributions $\mathcal{P}$. The probability model $P_{a,b}$ generates some random variable $x \in \mathcal{X}$ representing the observed data. Let $T(x)$ be a test of a null hypothesis $H_0$, e.g., $H_0 : (a, b) \in (\mathcal{A} \times \mathcal{B})_0 \subset \mathcal{A} \times \mathcal{B}$.

**Definition 0.1** (Identification Region)**.** The identification region $\text{Id}_{a,b}(P)$ is the pre-image of the model at a distribution $P \in \mathcal{P}$:

$$\text{Id}_{a,b}(P) = \{(a, b) \in \mathcal{A} \times \mathcal{B} : P_{a,b} = P\}.$$

**Definition 0.2** (Marginal Identification Region)**.** The marginal identification region of $b$ denoted $\text{Id}_b(P)$ at a model $P \in \mathcal{P}$ is

$$\text{Id}_b(P) = \{b \in \mathcal{B} : \exists\, a \in \mathcal{A}, P_{a,b} = P\}.$$

**Definition 0.3** (Marginal Identification)**.** The parameter $b$ is marginally unidentified at $P$ if $\text{Id}_b(P) = \mathcal{B}$. The parameter $b$ is marginally identified at $P$ if $\text{Id}_b(P)$ is a singleton set.

**Definition 0.4** (Level-$\alpha$ Hypothesis Test)**.** The test $T(x)$ of a null hypothesis $H_0 : (a, b) \in (\mathcal{A} \times \mathcal{B})_0$ is a level-$\alpha$ test if

$$\sup_{(a,b)\in(\mathcal{A}\times\mathcal{B})_0} \mathbb{E}_{P_{a,b}}[T(x)] \leq \alpha.$$

**Definition 0.5** (Trivial Hypothesis Test)**.** A test $T(x)$ of the null hypothesis $H_0 : (a, b) \in$

$(\mathcal{A} \times \mathcal{B})_0$ is a trivial test if it is level-$\alpha$ and

$$\sup_{(a,b)\in\mathcal{A}\times\mathcal{B}} \mathbb{E}_{P_{a,b}}[T(x)] \leq \alpha.$$

Marginal identifiability means that with sufficient samples from the model we can learn the exact values of the marginal parameter (e.g., $b$). Marginal unidentifiability means that no matter how many samples are observed from the model, we cannot learn anything about the marginal parameter (e.g., $b$). A trivial hypothesis test is no better than the test that just randomly rejects the null hypothesis with probability $\alpha$, regardless of the observed data. A trivial hypothesis test is therefore not useful in practice.

##### 0.1.1 Extended SRI Notation

The main text provided a brief summary of SRI theory and notation. Here we introduce additional concepts from SRI Theory [1] that we will use to establish theory for interval nulls.

As introduced in the main text, SRI models observed data $Y$ as an imperfect measurement from an underlying system $W$. More formally, SRI defines this measurement relationship probabilistically: $Y$ is generated by a probability model $P_{W^{\parallel},W^{\perp}}(Y)$ which is a single probability distribution out of a set of distributions $\mathcal{P}$. More generally, this is one distribution within the image of the model $P_{W^{\parallel},W^{\perp}}$ which is a mapping from the set of all possible systems $\mathcal{W}$. For brevity, we only consider positive-valued continuous systems $\mathcal{W} = \mathbb{R}^{D\times N+}$. The notion that $Y$ provides information about $W^{\parallel}$ but lacks information about $W^{\perp}$ is framed in terms of identifiability: within this model $W^{\parallel}$ is marginally identified whereas $W^{\perp}$ is marginally unidentified.

In SRI, the target estimand $\theta$ is a function of $W^{\parallel}$ and $W^{\perp}$. SRI also introduces a nuisance function $\lambda(W^{\parallel}, W^{\perp})$ such that there exists at least one $\lambda(W^{\parallel}, W^{\perp})$ that ensures that the following concatenated function $f$ is invertible:

$$(\theta, \lambda) = f(W^{\parallel}, W^{\perp}) = (\theta(W^{\parallel}, W^{\perp}), \lambda(W^{\parallel}, W^{\perp})).$$

This allows us to discuss $W^{\|}, W^{\perp}$ as a function of $\theta$: $(W^{\|}, W^{\perp}) = f^{-1}(\theta, \lambda)$. In turn we can reparameterize the measurement model as $P_{W^{\|},W^{\perp}} = P_{f^{-1}(\theta,\lambda)}$. SRI is defined as the situation where $\theta$ is marginally unidentified within this reparameterized model [1]. For brevity we omit the nuisance function $\lambda$ and the concatenated function $f$ from our notation and will simply write $P_\theta$ when discussing the reparameterized model.

Next, we use this reparameterized model to establish our theory of interval nulls.

##### 0.1.2 Theory of Interval Nulls for DA/DE in SRI

DE/DA centers on the LFC target estimand $\theta_d = \theta_d^{\|} + \theta^{\perp}$ defined in the main text. Consider a null hypothesis $H_0 : \theta_d = 0$ which can alternatively be written as $H_0 : \theta_d^{\|} + \theta^{\perp} = 0$. Consider an interval assumption $\theta^{\perp} \in [\tilde{\theta}^{\perp} + \epsilon_l^{\perp}, \tilde{\theta}^{\perp} + \epsilon_u^{\perp}]$. We define *the interval null* corresponding to this interval assumption as $H_0^* : \theta_d^{\|} + \tilde{\theta}^{\perp} \in [-\epsilon_u^{\perp}, -\epsilon_l^{\perp}]$.

**Theorem 1.** *Suppose $T(Y)$ is a non-trivial level-$\alpha$ test of the interval null hypothesis $H_0^*$ : $\theta_d^{\|} + \tilde{\theta}^{\perp} \in [-\epsilon_u^{\perp}, -\epsilon_l^{\perp}]$. Then, $T(Y)$ controls the false positive rate for, and is a non-trivial test of, $H_0 : \theta_d = 0$ if $\theta^{\perp} \in [\tilde{\theta}^{\perp} + \epsilon_l^{\perp}, \tilde{\theta}^{\perp} + \theta_u^{\perp}]$.*

*Proof.* We first prove that, if $\theta^{\perp} \in [\hat{\theta}^{\perp} + \epsilon_l^{\perp}, \theta^{\perp} + \theta_u^{\perp}]$, then $T(Y)$ controls the false positive rate for $H_0$. We then prove that it is non-trivial test of $H_0$.

By contradiction, suppose $T(Y)$ is a level-$\alpha$ test of $H_0^*$ but does not control the false positive rate for $H_0$ when $\theta^{\perp} \in [\hat{\theta}^{\perp} + \epsilon_l^{\perp}, \theta^{\perp} + \theta_u^{\perp}]$. Then,

$$\sup_{\{(\theta_d^{\|},\theta^{\perp}):\theta^{\perp}\in[\hat{\theta}^{\perp}+\epsilon_l^{\perp},\theta^{\perp}+\theta_u^{\perp}],\theta_d^{\|}+\theta^{\perp}=0\}} \mathbb{E}_{P_{\theta_d^{\|},\theta^{\perp}}}[T(Y)] > \alpha.$$

This is equivalent to

$$\sup_{\{(\theta_d^{\|},\theta^{\perp}):\theta^{\perp}\in[\hat{\theta}^{\perp}+\epsilon_l^{\perp},\theta^{\perp}+\theta_u^{\perp}],\theta_d^{\|}+\hat{\theta}^{\perp}\in[-\epsilon_u^{\perp},-\epsilon_l^{\perp}]\}} \mathbb{E}_{P_{\theta_d^{\|},\theta^{\perp}}}[T(Y)] > \alpha.$$

However, since $T(Y)$ is a level-$\alpha$ test of $H_0^* : \theta_d^{\|} + \hat{\theta}^{\perp} \in [-\epsilon_u^{\perp}, -\epsilon_l^{\perp}]$ it follows that

$$\sup_{\{(\theta_d^{\|},\theta^{\perp}):\theta^{\perp}\in[\hat{\theta}^{\perp}+\epsilon_l^{\perp},\theta^{\perp}+\theta_u^{\perp}],\theta_d^{\|}+\hat{\theta}^{\perp}\in[-\epsilon_u^{\perp},-\epsilon_l^{\perp}]\}} \mathbb{E}_{P_{\theta_d^{\|},\theta^{\perp}}}[T(Y)] \leq \alpha.$$

Hence there is a contradiction and $T(Y)$ must control the false positive rate for $H_0$ if it is a level-$\alpha$ test of $H_0^*$ and $\theta^\perp \in [\hat{\theta}^\perp + \epsilon_l^\perp, \theta^\perp + \theta_u^\perp]$.

Again by contraction, suppose $T(Y)$ is a non-trivial test of $H_0^*$ but is a trivial test of $H_0$. Then, the fact that it is a trivial test of $H_0$ implies that

$$\sup_{(\theta_d^\parallel, \theta^\perp) \in \mathbb{R}^2} \mathbb{E}_{P_{\theta_d^\parallel, \theta^\perp}}[T(Y)] \leq \alpha.$$

If $T(Y)$ is a non-trivial test of $H_0^*$ then there must be at least one element $(\theta_d^\parallel, \theta^\perp) \in \mathbb{R}^2$ at which $\mathbb{E}_{P_{\theta_d^\parallel, \theta^\perp}}[T(Y)] > \alpha$. This is a contradiction, therefore $T(Y)$ must be a non-trivial test of $H_0$ if it is a non-trivial test of $H_0^\star$. □

#### Derivation of Equivalence to Difference in Means of Log-transformed, Normalized Relative Abundances

In the main text we claim the equivalence

$$\theta_d^\parallel + \tilde{\theta}^\perp = \operatorname*{mean}_{n:x_n=1}(\log \tilde{W}_{dn}) - \operatorname*{mean}_{n:x_n=0}(\log \tilde{W}_{dn})$$

where $\tilde{W}^\perp = 1/\phi(W_{\cdot n}^\parallel)$ and therefore $\tilde{W}_{dn} = W_{dn}^\parallel / \phi(W_{\cdot n}^\parallel)$. The derivation of this equivalence may be written as follows:

$$\begin{aligned}
&\operatorname*{mean}_{n:x_n=1}(\log \tilde{W}_{dn}) - \operatorname*{mean}_{n:x_n=0}(\log \tilde{W}_{dn}) \\
&\quad = \operatorname*{mean}_{n:x_n=1}\left(\frac{\log W_{dn}^\parallel}{\phi(W_{\cdot n}^\parallel)}\right) - \operatorname*{mean}_{n:x_n=0}\left(\log \frac{W_{dn}^\parallel}{\phi(W_{\cdot n}^\parallel)}\right) \\
&\quad = \operatorname*{mean}_{n:x_n=1}\left(\log W_{dn}^\parallel + \log \frac{1}{\phi(W_{\cdot n}^\parallel)}\right) - \operatorname*{mean}_{n:x_n=0}\left(\log W_{dn}^\parallel + \log \frac{1}{\phi(W_{\cdot n}^\parallel)}\right) \\
&\quad = \operatorname*{mean}_{n:x_n=1}\left(\log W_{dn}^\parallel\right) - \operatorname*{mean}_{n:x_n=0}\left(\log W_{dn}^\parallel\right) + \operatorname*{mean}_{n:x_n=1}\left(\log \frac{1}{\phi(W_{\cdot n}^\parallel)}\right) - \operatorname*{mean}_{n:x_n=0}\left(\log \frac{1}{\phi(W_{\cdot n}^\parallel)}\right) \\
&\quad = \theta_d^\parallel + \tilde{\theta}^\perp.
\end{aligned}$$

#### Connection Between the CTT and Sensitivity Analyses

In the main text we considered the null hypothesis of the form $H_0 : \theta_d = 0$. By assuming $\theta^\perp \in [\tilde{\theta}^\perp + \epsilon_l^\perp, \tilde{\theta}^\perp + \epsilon_u^\perp]$ we derived the interval null hypothesis $H_0^* : \theta_d^{||} + \tilde{\theta}^\perp =\in [-\epsilon_u^\perp, -\epsilon_l^\perp]$. We then derived the Composite T-Test (CTT) by its p-value as follows. Let $p_-$ be the p-value from a one-sided test of $H_{0-} : \theta_d^{||} + \tilde{\theta}^\perp \leq -\epsilon_u^\perp$, and $p_+$ the p-value from a test of $H_{0+} : \theta_d^{||} + \tilde{\theta}^\perp \geq -\epsilon_l^\perp$. The CTT p-value for for $H_0^* : \theta_d^{||} + \tilde{\theta}^\perp \in [-\epsilon_u^\perp, -\epsilon_l^\perp]$ can then be constructed as:

$$p^{\text{CTT}} = \min\{1, 2p_-, 2p_+\}.$$

However, rather than apply the CTT to the interval null hypothesis, consider instead a sensitivity analysis where we test the null hypothesis $H_0 : \theta_d^{||} + \tilde{\theta}^\perp + \epsilon^\perp = 0$ where $\epsilon^\perp$ is error in $\tilde{\theta}^\perp$. For any one $\epsilon^\perp$, a simple two-sided Student's t-test of this null hypothesis could be performed to derive a p-value $p_{\epsilon^\perp}$. We can then perform a sensitivity analysis by visualizing how $p_{\epsilon^\perp}$ changes as a function of $\epsilon^\perp$.

However, note that the interval assumption $\theta^\perp \in [\tilde{\theta}^\perp + \epsilon_l^\perp, \tilde{\theta}^\perp + \epsilon_u^\perp]$ is equivalent to assuming $\epsilon^\perp \in [\epsilon_l^\perp, \epsilon_u^\perp]$ for the null hypothesis $H_0 : \theta_d^{||} + \tilde{\theta}^\perp + \epsilon^\perp = 0$. We can then use an interval assumptions to turn a sensitivity analysis into a *sensitivity test* defined by the p-value:

$$p^{\text{SENS}} = \sup_{\epsilon^\perp \in [\epsilon_l^\perp, \epsilon_u^\perp]} p_{\epsilon^\perp}.$$

The connection between our work and sensitivity analyses is that the sensitivity test and the Composite T-Test (CTT) of the interval null hypothesis $H_0^*$ are equivalent statistical tests.

We prove that these tests are equivalent by proving $p^{\text{CTT}} = p^{\text{SENS}}$ as follows. In the sensitivity test, we test the null hypothesis $H_0 : \theta_d^{||} + \tilde{\theta}^\perp + \epsilon^\perp = 0$ by applying a two-sided t-test for each individual $\epsilon^\perp \in [\epsilon_l^\perp, \epsilon_u^\perp]$. Note that this is equivalent to testing $H_0 : \theta_d^{||} + \tilde{\theta}^\perp = -\epsilon^\perp$ for all $\epsilon^\perp \in [\epsilon_l^\perp, \epsilon_u^\perp]$. Consider the case that $\theta_d^{||} + \tilde{\theta}^\perp \in [-\epsilon_u^\perp, -\epsilon_l^\perp]$, the p-value $p^{\text{SENS}}$ in this case must always be 1 and the p-value $p^{\text{CTT}}$ must also always be 1. If instead, $\theta_d^{||} + \tilde{\theta}^\perp < -\epsilon_u^\perp$, the p-value $p^{\text{CTT}}$ will be $2 \times p_-$ which is equivalent to the p-value for a two-sided t-test of the null hypothesis $H_0 : \theta_d^{||} + \tilde{\theta}^\perp = -\epsilon_u^\perp$. The p-value $p^{\text{SENS}}$ will also be equivalent to a two-sided t-test of the null hypothesis $H_0 : \theta_d^{\|} + \tilde{\theta}^{\perp} = -\epsilon_u^{\perp}$. This same logic may be applied to the case that $\theta_d^{\|} + \tilde{\theta}^{\perp} > -\epsilon_l^{\perp}$. This proves that $p^{\mathrm{CTT}} = p^{\mathrm{SENS}}$ in all cases.

We propose the CTT rather than the sensitivity test in the main text because the CTT can be defined most clearly as a test of the interval null hypothesis. The interval null hypothesis is a more flexible framework for defining statistical tests. Specifically, the interval null hypothesis allows us to apply the Generalized T-Test (GTT) to this problem, which is more powerful than the CTT or sensitivity test [2].

#### Deriving Confidence Intervals from Interval Null Hypothesis Tests

Here we derive confidence intervals by inverting the CTT and GTT tests. Given an interval assumption of the form $\theta^{\perp} \in [\tilde{\theta}^{\perp} + \epsilon_l^{\perp}, \tilde{\theta}^{\perp} + \epsilon_u^{\perp}]$, the confidence intervals derived by inverting both the CTT and GTT satisfy

$$\inf_{(\theta_d^{\|}, \theta^{\perp}) \in \mathbb{R}^2} P_{\theta_d^{\|}, \theta^{\perp}} \left( \theta^{\|} + \tilde{\theta}^{\perp} - \epsilon^{\perp} \in C(Y) \right) \geq 1 - \alpha, \, \forall \epsilon^{\perp} \in [-\epsilon_u^{\perp}. - \epsilon_l^{\perp}].$$

However, using the relationship $\theta_d = \theta_d^{\|} + \theta^{\perp}$, this guarantee may be equivalently expressed as

$$\inf_{\{(\theta_d^{\|}, \theta^{\perp}) \in \mathbb{R}^2 : \theta^{\perp} \in [\tilde{\theta}^{\perp} + \epsilon_l^{\perp}, \tilde{\theta}^{\perp} + \epsilon_u^{\perp}]\}} P_{\theta_d^{\|}, \theta^{\perp}} \left( \theta_d \in C(Y) \right) \geq 1 - \alpha.$$

Therefore, the following confidence intervals guarantee coverage of the LFC $\theta_d$ with probability $1 - \alpha$, but only if the interval assumption $\theta^{\perp} \in [\tilde{\theta}^{\perp} + \epsilon_l^{\perp}, \tilde{\theta}^{\perp} + \epsilon_u^{\perp}]$ is true.

Let $C_{\epsilon^{\perp}}(Y)$ be the standard $1 - \alpha$ confidence interval derived by inverting a two-sided Student's t-test of the null hypothesis $H_0 : \theta^{\|} + \tilde{\theta}^{\perp} = \epsilon^{\perp}$. Then the confidence interval corresponding to the CTT is defined as

$$C^{\mathrm{CTT}}(Y) = \bigcup_{\epsilon^{\perp} \in [-\epsilon_u^{\perp}, -\epsilon_l^{\perp}]} C_{\epsilon^{\perp}}(Y).$$

Following Mehring [2] and using the notation from the methods section in the main text, the GTT may be equivalently defined by its crirical value $c$, which solves the equation

$$1 + F_v((-c - \psi_2')/\sigma_{\mathrm{SE}}) - F_v((c - \psi_2')/\sigma_{\mathrm{SE}}) = \alpha.$$

The GTT test can then be defined by the decision function that rejects $H_0'$ if the estimated difference in means $\hat{\mu}_\Delta$ satisfies $\hat{\mu}_\Delta < -c$ or $\hat{\mu}_\Delta > c$. Using $c$, the confidence interval corresponding to the GTT can then be written as

$$C^{\mathrm{GTT}}(x, y) = [\hat{\mu}_\Delta - \bar{\psi} - c, \hat{\mu}_\Delta - \bar{\psi} + c].$$

For DE/DA, $\bar{\psi} = (-\epsilon_u^\perp - \epsilon_l^\perp)/2$, $\hat{\mu}_\Delta$ represents the estimate of $\theta_d^{\|} + \tilde{\theta}^\perp$, and $\psi_2' = -\epsilon_l^\perp - \bar{\psi}$.

### Supplementary Figures

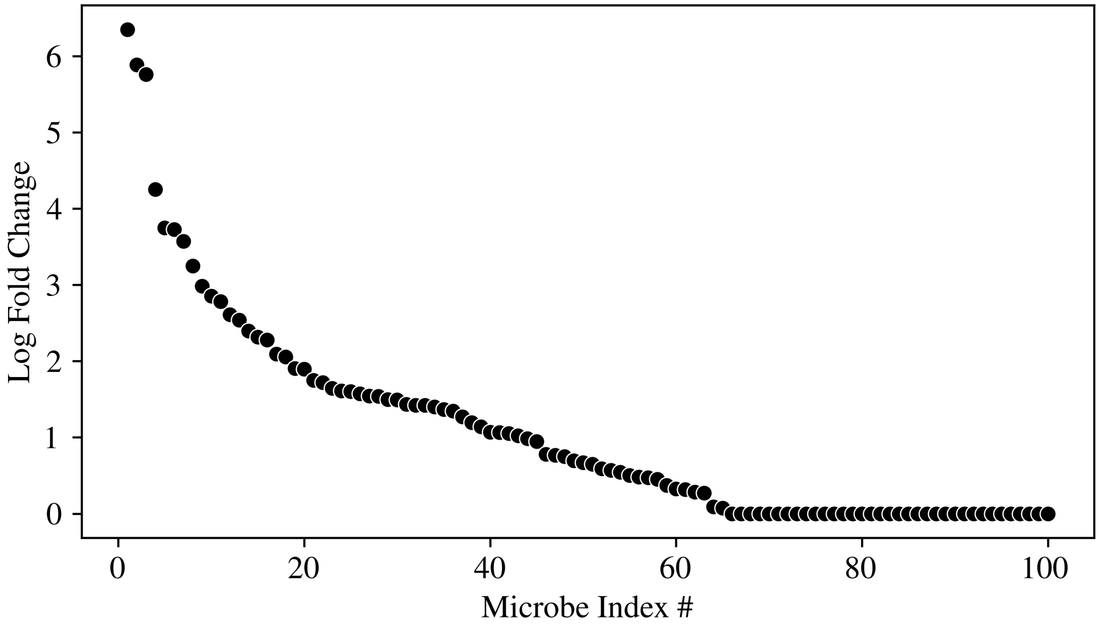


**Figure S1: The LFC of the microbes used in the SpraseDOSSA2 simulation presented in Figure 2 of the main text.**

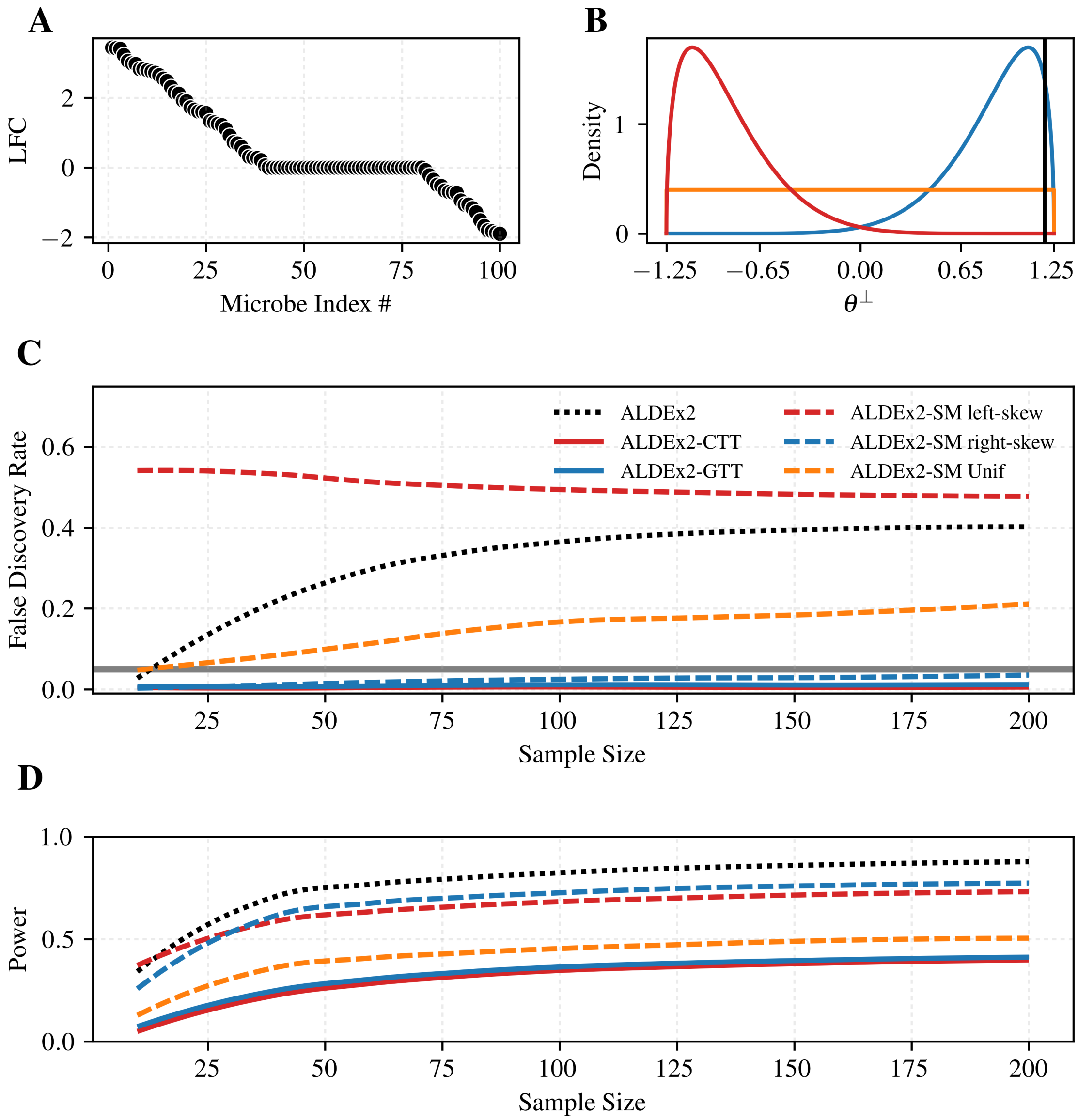


**Figure S2: Interval assumptions guarantee control of the false discovery rate if the interval covers $\theta^{\perp}$, unlike scale models. A.** The LFC of 100 microbes which were used to simulate absolute abundances from a multivariate normal distribution. **B.** Three scale models used in an ALDEx2 analysis all defined over the interval $[-1.25, 1.25]$ including a uniform distribution (orange), and left- and right-skewed distributions (see *Methods*; red and blue respectively). The true $\theta^{\perp} = 1.22$ is shown as a vertical black line. **C.** The average false discovery rate over 100 independent simulations at each sample size. Methods used include ALDEx2 (CLR normalization), ALDEx2 with each scale model, and ALDEx2-CTT and ALDEx2-GTT using the interval assumption $\theta^{\perp} \in [-1.25, 1.25]$ matching the support of the scale models. Only ALDEx2-CTT, ALDEx2-GTT, and ALDEx2 with the right-skew scale model controlled the false discovery rate at the desired level 0.05 (gray horizontal line). **D.** The average power of each method at each sample size over the same 100 independent simulations as in part C.
